# Development of an Application Ontology for Beta Cell Genomics Based On the Ontology for Biomedical Investigations

**DOI:** 10.1101/2025.08.18.670933

**Authors:** Jie Zheng, Elisabetta Manduchi, Christian J. Stoeckert

## Abstract

The development process for the beta cell genomics application ontology (BCGO) is described. This process should be generally applicable and consists of integration of a subset of reference ontologies. A key element is use of the Ontology for Biomedical Investigation (OBI) as an ontology framework. Another element is enriching ontologies using existing patterns when needed. The ontology is validated in three aspects based on our needs including data annotation, queries and automated classification. The BCGO is available on: http://purl.obolibrary.org/obo/bcgo.owl.

## 1 INTRODUCTION

The Beta Cell Genomics database is a functional genomics resource focused on pancreatic beta cell research (http://genomics.betacell.org/gbco/) and contains 128 public and private studies (in v4.11). It supports the National Institute of Diabetes and Digestive and Kidney Diseases Beta Cell Biology Consortium (http://www.betacell.org) in its mission to advance understanding of pancreatic islet development and function, with the goal of developing therapies to correct the loss of beta cell mass in diabetes. Much of the current research focuses on producing functional beta cells either from stem cells or reprogramming mature cells of other types such as exocrine cells (Borowiak and Melton, 2009). One challenge is establishing criteria (both biological features and genetic signatures) that can be used to determine whether a reprogrammed or differentiated cell is a functional beta cell. It has been demonstrated that semantic descriptions can enhance queries and facilitate knowledge discovery (*e.g*. automated classification) using reasoners based on computable definitions (Askenazi and Linial, 2011, Köhler *et al*., 2012, Malone *et al*., 2010, Meehan *et al*., 2011). This approach can be applied to effectively find related studies and data on particular types of cells and model systems in the Beta Cell Genomics database and learn about cells cultured or reprogrammed to achieve a desired cell fate or phenotype. Moreover, an ontology with enough granularity for beta cell studies could be used to support reasoning and cover both biological and experimental aspects of functional genomics studies.

The Open Biological and Biomedical Ontologies (OBO) Foundry established a set of principles for developing orthogonal interoperable ontologies in biomedical domains aiming to facilitate ontology integration (Smith *et al*., 2007). OBO Foundry (candidate) ontologies are built on the basis of a common top-level ontology, the Basic Formal Ontology (BFO), and use a common set of relations, primarily defined in the Relation Ontology (RO) (Smith *et al*., 2005). Each OBO Foundry reference ontology covers a specific domain. For example, the Cell Ontology (CL) defines native cell types (Meehan *et al*., 2011), the Cell Line Ontology (CLO) represents *in vitro* cell lines (Sarntivijai *et al*., 2011), the Gene Ontology (GO) focuses on cellular component, molecular function, and biological process related to genes and gene products (Ashburner *et al*., 2000), and the Uber anatomy ontology (UBERON) is used for cross-species anatomy annotation (Mungall *et al*., 2012).

The Ontology for Biomedical Investigations (OBI) has been developed following OBO Foundry principles for supporting consistent representation of biological and clinical investigations including functional genomics studies (Brinkman *et al*., 2010). BFO is used as the top-level ontology and the Information Artifact Ontology (IAO) is used to represent ontology metadata and information such as data, investigation design and textual entities. OBI describes all aspects of an investigation including biological materials, assays, protocols, generated data and type of analysis applied to the data. OBI contains classes (*e.g*. cells, gross anatomical parts, biological processes) that are important for modeling biomedical investigations but are out of its scope. These classes can be used to connect OBI with other OBO Foundry reference ontologies, such as CL, UBERON, and GO, and serve as the parent of referenced external terms.

There is no single existing OBO Foundry ontology that can currently meet the prescribed needs for Beta Cell Genomics. The use of multiple ontologies to annotate Beta Cell studies brings in unnecessary complexity since only small subsets of reference ontologies are used and not all defined logic axioms are useful. Malone *et al* have created the application ontology, Experimental Factor Ontology (EFO), by reusing reference ontologies when available and enriching the ontology with additional axioms when needed (Malone *et al*., 2010). With well-established ontologies now available in the OBO community, we are able to adapt the EFO approach to build the Beta Cell Genomics Ontology (BCGO) by integration of OBO Foundry ontologies based on the OBI framework and reusing existing ontology design patterns to enrich the ontology. OBI is used as the basis to create the BCGO because it represents all aspects of an experiment and can integrate with other OBO ontologies through links to external resources (http://purl.obolibrary.org/obo/obi).

## 2 METHODS

The BCGO was built based on the integration of multiple OBO Foundry ontologies and then enriched with terms specific to Beta Cell Genomics research. Therefore, we needed first to identify which ontologies are relevant to the Beta Cell Genomics studies. Terms of interest were then extracted from the relevant ontologies and integrated. Details of the methodology for these steps are described below.

### 2.1 Identification of OBO Foundry ontologies

The Beta Cell Genomics database contains studies annotated using multiple controlled vocabularies and ontologies including the MGED Ontology (MO) (Whetzel *et al*., 2006). The terms used in the annotation were extracted from the Beta Cell Genomics database and mapped to the OBO Foundry ontologies. The terms were first checked using the MO to OBI mapping list on: https://github.com/obi-bcgo/bcgo/blob/master/doc/mapping%20results/MO2OBI_Mappings.xlsx.

Terms not found in the MO to OBI mapping were mapped to OBO Foundry ontologies using the BioPortal annotator web services (Jonquet *et al*., 2009). The annotator service can accurately (>95%) tag text with ontology terms. However, ontologies in the annotator might not be the latest version since these need to go through an indexing process before being added to the annotator. Unmapped terms were then searched in the ontologies using the BioPortal search web services (Whetzel *et al*., 2011). Both annotator and search results were reviewed manually. The search results required more manual review effort since that search provides both partial and exact matches of input text.

### 2.2 Application ontology development

The W3C standard Web Ontology Language Description Logic (OWL-DL) was used to implement the BCGO ontology to provide rich semantics and support for automated reasoning and inferences. Protégé 4.2 was used for editing the ontology and Hermit 1.3.6 used for reasoning and consistency checking.

Development of the application ontology includes three steps (shown in Figure 1):

**Fig. 1.**
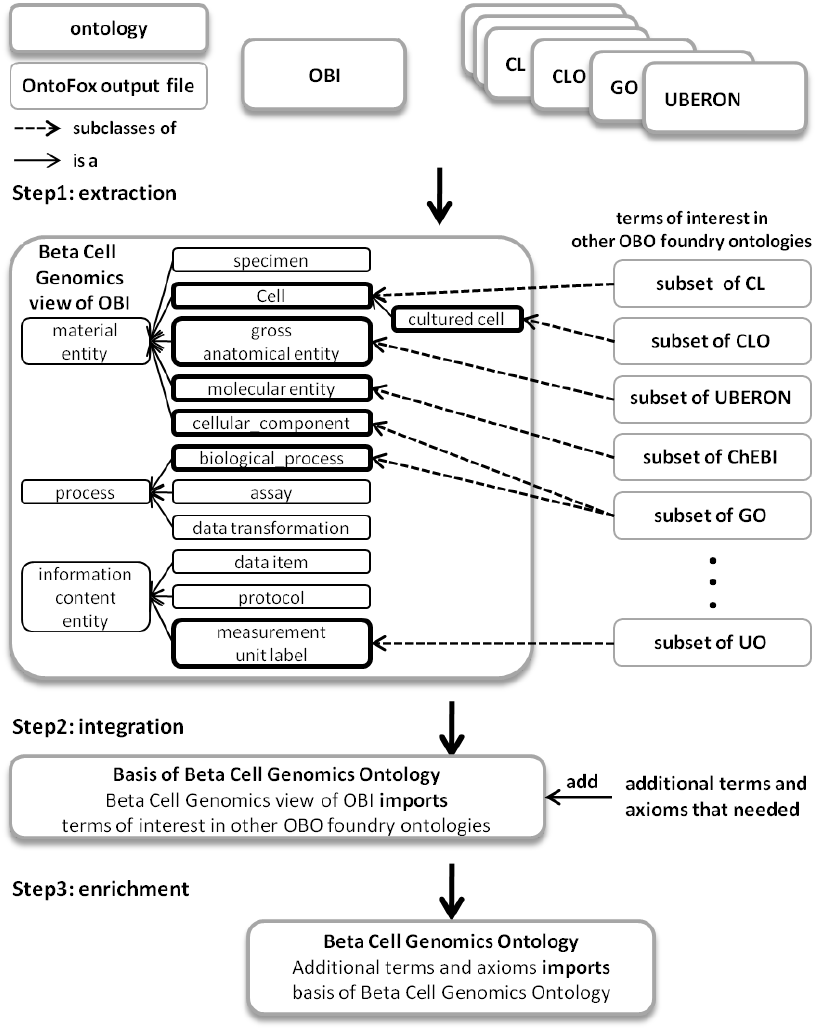
Main steps of BCGO development: (1) ontology extraction (2) integration of ontology subsets and (3) ontology enrichment. ChEBI stands for Chemical Entities of Biological Interest ontology and UO for Units of Measurement ontology.

1. Extraction: retrieve terms of interest from ontologies including OBI;
2. Integration: integrate ontology terms retrieved from various ontologies into the retrieved OBI subset;
3. Enrichment: enrich the integrated subset ontology by adding additional terms and axioms where needed using existing ontology design patterns if available.

The application ontology consists of subsets of various ontologies and additional terms and logical restrictions needed for our specific application. A layered ontology modules approach (Torniai *et al*., 2011) was used to decouple different components mainly for maintenance purposes. Each component was kept in an individual OWL file. The OWL import mechanism was used to group different components together and construct the application ontology. If retrieved labels, definitions or logical axioms are updated in a source ontology, they will only need to be updated in that ontology component in the application ontology. The layered ontology modules approach provides flexibility in synchronization with source ontologies and facilitates parallel development.

The BFO was used as the top-level ontology and imported as a whole into the BCGO. The meta-data schema was implemented as OWL annotation properties defined in the IAO and widely used in the OWL format ontologies, such as OBI, CL, CLO and GO.

#### 2.2.1 Ontology Extraction

Since the BCGO is built on the framework of OBI, the retrieved subset of OBI provides an upper level hierarchical structure covering all aspects of an investigation. The generic OBI terms for a functional genomics investigation that were also used in the Beta Cell Genomics database were retrieved from OBI using Ontodog (Zheng *et al*., 2012). Ontodog is a web-based tool that can retrieve a selected set of terms from a source ontology including all relevant terms and logical axioms and retain reasoning consistency. The output file is in the OWL format.

The terms needed from other relevant OBO Foundry ontologies were retrieved using OntoFox (Xiang *et al*., 2010) with the OBI view created with Ontodog serving as the target ontology. OntoFox is a web-based system that allows users to retrieve a set of terms in ontologies and specify the super-classes in the target ontology for reusing the external terms in the target ontology. The output of OntoFox is an OWL format file that can be imported into a target ontology of the same format. OntoFox supports various approaches to fetch terms from an ontology. Three options were considered:

1. Retrieve minimum information of a term from the source ontology that allows the term to be referenced in the target ontology. Based on the Minimum Information to Reference an External Ontology Term (MIREOT) guideline, one needs to specify the URI of the term, the IRI of the source ontology and the term’s parent URI in the target ontology.
2. Retrieve a set of terms from the source ontology using the “includeComputedIntermediates” setting which will fetch the terms of interest with minimal intermediate ontology terms between specified upper and lower terms. The extracted terms can keep the hierarchy organization excluding other logical axioms defined in the source ontology.
3. Retrieve a set of terms from the source ontology using a SPARQL-based related term retrieval algorithm by choosing the “includeAllIntermediates” and “includeAllAxiomsRecursively” settings. The retrieved subset ontology includes all relevant axioms and terms used in the axioms.

All retrieval options will fetch the label and definition of terms from the source ontologies. Option 1 can be done manually too.

Our choice of the extraction method depended on the desired application. When logical axioms were needed for the application, option 3 was used. Option 2 was used for keeping the hierarchical structure of the terms in the source ontologies. Generally, option 1 was used when few terms (generally 5 or less) in an ontology were referenced in the application ontology.

OntoFox can accept input data from a local text file via uploading. The input data file is reusable and used to synchronize with source ontologies. To support the implementation of layered ontology modules, a separate OntoFox input data file was created for each reference ontology to generate corresponding output OWL files that contain the terms of interest from each source ontology.

#### 2.2.2 Ontologies integration

OBO Foundry reference ontologies may contain terms referencing external resources. For example, the CL ontology uses GO, the Protein ontology (PR), and UBERON to define a cell by specifying its function, expressed protein on the membrane and the anatomical structure to which it belongs. Using the Ontodog and OntoFox option 3 approach to extract a subset of an ontology may result in inclusion of terms defined in external ontologies. For terms in the source ontology imported from external ontologies, both textual and logical definitions in the retrieved subset were removed to avoid possible conflicts as they may have diverged from the original definitions. Terms of interest from the same source ontology were combined and retrieved together.

The output files of OntoFox were imported directly into the target ontology using *owl:imports* statement. Consistency of the integrated ontology was confirmed using the ontology reasoner.

#### 2.2.3 Ontology enrichment

Terms unavailable in the OBO Foundry ontologies will be submitted to corresponding reference ontologies according to their scopes and defined using ontology design patterns when available. For example, data analysis terms will be submitted to OBI. The input and output data, analysis objective, and algorithm used (if applicable) will be specified based on the OBI developed pattern.

Needed terms not belonging to the scope of any existing reference ontologies as well as desired but unavailable logical restrictions will be implemented in the BCGO ontology.

## 3 RESULTS

### 3.1 Identification of OBO Foundry ontologies

Out of 852 terms used in the Beta Cell Genomics database, 644 terms were matched to 543 ontology terms defined in 24 various ontologies including BFO (1 term), IAO (2 terms), 19 OBO Foundry reference ontologies, 3 OBO Foundry application ontologies, BRENDA tissue/enzyme source (BTO), EFO, and eagle-i research resource ontology (ERO) (shown in Table 1, Matched Terms column). The matched terms primarily belong to reference ontologies, OBI, UBERON, UO (Units of measurement), CL, PATO (Phenotypic quality), ChEBI and CLO. The results show that only small portions of each ontology were needed for the Beta Cell Genomics database.

While it is desirable to build an application ontology just from reference ontologies to avoid overlap in terms, it was necessary to include terms from other application ontologies. Forty EFO terms (mainly developmental stages and diseases) and two ERO terms (1 planned process and 1 assay) not available in reference ontologies were used. All ontologies listed in the Table 1 Ontology column were used to build the BCGO.

**Table 1.** Mapping and extraction of terms. Ontology refers to the namespace of OBO Foundry ontologies. CARO: Common Anatomy Reference Ontology; EnVO: Environment Ontology; FMA: Foundational Model of Anatomy; GAZ: Gazetteer; MP: Mammalian Phenotype; NCBITaxon: NCBI organismal classification; OGMS: Ontology for General Medical Science; RS: Rat Strain ontology: SWO: Software Ontology. Other namespaces and their associated ontology are described in the text. Version refers to the ontology version number or release date used for mapping and extraction. N/A indicates not available. Total Classes are the total number of classes with namespace ID associated with each ontology. Matched Terms are the number of Beta Cell Genomics database terms matching the ontology terms. Relevant Classes are the number of classes needed for the BCGO including those used in the database and in logic axioms. Extracted Classes are the number of classes retrieved from the listed ontology. Extraction Method indicates whether Ontodog or OntoFox (option 1, 2 or 3) were used for ontology extraction. ^*^: application ontology

| Ontology | Version | Total Classes | Matched Terms | Relevant Classes | Extracted Classes | Extraction Method |
| --- | --- | --- | --- | --- | --- | --- |
| OBI | 2012-07-01 | 2042 | 200 | 201 | 430 | ontodog |
| BTO* | 12/20/2012 | 5391 | 2 | 2 | 2 | option 1 |
| CARO | N/A | 50 | 1 | 1 | 1 | option 1 |
| EnVO | 2013-01-08 | 1557 | 1 | 2 | 2 | option 1 |
| ERO* | 2012-10-03 | 1579 | 2 | 2 | 2 | option 1 |
| FMA | 3.1 | 83281 | 1 | 1 | 1 | option 1 |
| GAZ | 1.512 | 518195 | 1 | 1 | 1 | option 1 |
| MP | 07/14/2012 | 9164 | 1 | 1 | 1 | option 1 |
| OGMS | 2011-09-20 | 81 | 3 | 3 | 3 | option 1 |
| RS | 1/14/2013 | 3361 | 1 | 1 | 1 | option 1 |
| SO | 11/1/2012 | 2151 | 1 | 5 | 5 | option 1 |
| SWO | 0.5 | 661 | 1 | 1 | 1 | option 1 |
| EFO* | 2.31 | 4057 | 40 | 40 | 65 | option 1,2 |
| ChEBI | 100 | 38901 | 12 | 14 | 62 | option 2 |
| CLO | 2.1.03 | 35436 | 11 | 11 | 19 | option 2 |
| GO | 2012-12-18 | 38747 | 2 | 80 | 164 | option 2 |
| NCBITaxon | 2013-01-24 | 981148 | 1 | 14 | 20 | option 2 |
| PR | 31.0. | 35488 | 1 | 55 | 59 | option 2 |
| UO | 2012-08-30 | 313 | 67 | 67 | 74 | option 2 |
| CL | 2013-01-31 | 2120 | 46 | 46 | 309 | option 3 |
| PATO | 01/09/2013 | 2426 | 19 | 33 | 70 | option 3 |
| UBERON | 2013-01-07 | 7318 | 126 | 141 | 1206 | option 3 |

### 3.2 Ontology extraction

A subset of OBI including IAO terms was extracted using Ontodog to generate the Beta Cell Genomics view of OBI. Additional terms of interest defined in OBO Foundry ontologies were extracted using different retrieval approaches as described in the Methods and indicated in Table 1.

The CL ontology represents cell type based on several aspects, including biological processes in which the cell participates, associated phenotypic features, and protein complexes found on the surface of the cell (Meehan *et al*., 2011). Computable definitions of cell types enable automated classification based on various aspects of cells. The CL also contains high level types of cell (such as *in vitro* cell) that are reused by the CLO. The CLO relates *in vitro* cell to the native cell defined in CL using the relation *derived from*. PATO provides computational definitions to represent qualities relative to normal. UBERON connects different anatomical parts, relates anatomical entities to specific developmental stages, and covers high level life cycle stages. All of these logical axioms are useful for issuing complicated queries and automated classification. Therefore, extraction option 3 was used to fetch CL, PATO, and UBERON terms using the OntoFox tool. Option 2 was used for the CLO term retrieval due to some known issues but option 3 will be used when these are fixed for the official release.

Many ChEBI, GO, NCBITaxon, PR, and UO terms were used in the database and needed for defining other ontology classes. Extraction option 2 was used to preserve the hierarchy in these ontologies as the hierarchy aids ontology term searches. Matched EFO terms for developmental stages were also retrieved using option 2. The rest of the matched terms were extracted and referenced using option 1.

### 3.3 Ontology integration

The subsets of OBO Foundry ontologies were integrated into the Beta Cell Genomics view of OBI using OWL imports. The root classes of each subset were aligned to the OBI outer core terms. The integrated ontology is the basis of the BCGO and enriched with needed terms and restrictions. There are multiple versions of BFO available. Most of the OBO Foundry ontologies used to construct the BCGO basis including CL, GO, OBI, PATO, PR, UBERON, and UO are using the BFO 2.0 pre-Graz version. Therefore, this version of BFO is used by the BCGO. The base BCGO is available at: https://raw.githubusercontent.com/obi-bcgo/bcgo/refs/heads/master/release/v0.1/bcgo.owl https://raw.githubusercontent.com/obi-bcgo/bcgo/refs/heads/master/release/v0.1/bcgo_basis.owl. It contains 2371 classes. Consistency of the ontology was confirmed by OWL reasoner Hermit 1.3.6.

### 3.4 Ontology enrichment

The 208 terms that could not be matched to OBO Foundry ontologies fall mainly in the scope of OBI (46 terms), CL (35 terms), CLO (23 terms) and UBERON (33 terms). These terms will be submitted to the corresponding reference ontology with both textual and logical definitions using existing ontology design patterns when available. For example, ‘Ngn3 null pancreatic progenitor cell’ can be textually defined as ‘a multi fate stem cell of embryonic pancreatic buds with no expression of neurogenin-3’ and logically defined (in OWL) as:

> *‘Ngn3-null pancreatic progenitor cell’*
>
> *subClassOf CL:’multi fate stem cell’* and
>
> *BFO:is_part_of* some *UBERON:’pancreatic bud’* and
>
> *CL:lacks_part* some *PR:neurogenin-3*.

The CL describes protein products related to a cell. Beta cell research is often at the gene expression level and many studies focus on finding genetic signatures of a cell. To represent gene expression information of a cell in the BCGO, we added axioms of the type:

> *RO:produces* some *(SO:transcript* and
>
> *(SO:translates_to* some *PR:protein))*

RO and SO (Sequence types and features Ontology; Mungall *et al*., 2011) were used to supplement BFO relations.

For example, two additional axioms were added to *CL: ‘progenitor cell of endocrine pancreas’* which has the textual definition “A multi-fate stem cell that is able to differentiate into the pancreas alpha, beta and delta endocrine cells. This cell type expresses neurogenin-3 and Isl-1”:

> *RO:produces* some *(SO:transcript* and
>
> *(SO:translates_to* some *PR: insulin gene enhancer protein ISL-1))*
>
> *RO:produces* some *(SO:transcript* and
>
> *(SO:translates_to* some *PR: neurogenin-3))*

Twenty eight unmatched terms are strains (mainly mouse strains). The RS is the only available OBO Foundry candidate ontology to represent strains. However, the RS does not have an ontology design pattern to define genotypes of a strain based on genetic background, altered genetic information, and how a strain is generated or maintained. In OBI, strain is defined based on how it is generated:

> *OBI:organism* and *(OBI:is_specified_output_of* some
>
> *(OBI:’planned process’* and
>
> *(OBI:achieves_planned_objective* some
>
> *OBI:’selective organism creation objective’)))*

This pattern was used to define strains in the BCGO. For example, mouse strain is defined as:

> *NCBITaxon:’mus musculus’* and *(OBI:is_specified_output_of*
>
> some *(OBI:’planned process’* and
>
> *(OBI:achieves_planned_objective* some
>
> *OBI:’selective organism creation objective’)))*

To enable query of finding genomics data based on the genetic background of strains, logical representation of the genetic background of a strain is needed. We added an axiom to define a strain in the BCGO:

> *BCGO:has_genetic_background* some
>
> *(OBI:’genetic population background information’* and
>
> *(IAO:’is about’* some
>
> *OBI:’selectively maintained organism’))*

### 3.5 Ontology validation

The BCGO is an application ontology created for the Beta Cell Genomics database and required to support the following applications:

1. Data annotation
2. Enable complicated queries, such as “find high throughput sequencing gene expression data in samples obtained from mouse strains with genetic background C57BL/6J during the embryo stage”, “find gene expression data of endocrine cells”, “find studies using cells which develop from either mesoderm or endoderm”.
3. Automated classification, such as identify the type of cell based on biological features and genetic signatures.

The BCGO ontology was evaluated by checking whether it could meet the above desired needs.

#### 3.5.1 Data annotation

Currently, the BCGO contains about 80% of the terms used in the database. Ongoing work is to include the rest of them, either by submission to the appropriate reference ontologies as described in section 3.4 or directly adding into the BCGO.

#### 3.5.2 Complex queries

The BCGO allows us to issue sophisticated queries of the types just described. The query “find high throughput sequencing gene expression data in pancreatic cells derived from mouse strain with genetic background C57BL/6J during embryo stage” converts to data generated from *‘sequencing assay’*, and specimen used for assay collected from any *‘mouse strain’* that *has_genetic_background* some *‘genetic population genetic background’ ‘is about’* some *‘C57BL/6J strain’* at developmental stage *‘embryo stage’*. “Find studies using cells which develop from either mesoderm or endoderm” will return studies on cells that developed from endoderm (hepatocyte, epithelial cell of pancreas, type B pancreatic cell, etc.) and cells that developed from mesoderm (muscle cell).

#### 3.5.3 Automated classification

In the CL, it has been demonstrated that cells can be classified based on different criteria, such as cellular functions and cell surface markers, implemented in computable definitions (Meehan *et al*., 2011). Adopting these same mechanisms, the BCGO may help define specific cell types such as a functional beta cell through genetic signatures. It has been reported that features of functional beta cells include glucose sensing, secreting insulin, and expression of certain transcriptional regulators such as MafA, Pdx1, etc. (Benitez *et al*., 2012). To establish criteria for a functional beta cell that can be implemented in OWL, ‘functional beta cell like cell’ will be added in the BCGO with computable definitions covering all known features of functional beta cell including:

> *CL:capable_of* some *GO:’detection of glucose’*
>
> *CL:capable_of* some *GO:’insulin secretion’*
>
> *RO:produces* some *(SO:transcript* and
>
> *(SO:translates_to* some *PR:’transcription factor MafA’))*
>
> *RO:produces* some *(SO:transcript* and
>
> *(SO:translates_to* some *PR:’pancreas/duodenum homeobox protein 1’))*

With an OWL reasoner, ‘type B pancreatic cell’ should be inferred as a ‘functional beta cell like cell’. However, ‘pancreatic bud insulin expressing cell’, ‘muscle cell’, and hepatocyte should not be inferred as a ‘functional beta cell like cell’. Unexpected inferred relationships indicate wrong or incomplete logic definitions of terms.

Genomics data of over 20 different kinds of cells including pancreatic beta cell, alpha cell, progenitor cell of endocrine pancreas, embryonic stem cell, muscle cell, hepatocyte are available in the Beta Cell Genomics database. Known biological features of various cells as well as associated gene expression information generated from genomics data analysis will be added to corresponding cell types in the BCGO and used to test whether the computable definition of ‘functional beta cell like cell’ is correct. When no unexpected inference occurs, necessary and sufficient conditions defining ‘functional beta cell like cell’ can be used to encode criteria for a functional beta cell.

## 4 DISCUSSION

A cross-domain application ontology has been developed based on the OBI framework and reusing existent reference ontologies and ontology design patterns. The approach used to develop the BCGO should be generally applicable when using interoperable source ontologies. Differences in this approach from that used for EFO development include reusing the high level structure defined in the OBI, logical axioms defined in multiple reference ontologies, and existing ontology design patterns. It should be noted that the latter were generally unavailable during the period of the EFO development.

The big challenge of the approach described here is integration of ontology modules from various resources. Although OBO Foundry policy aims to facilitate this process, conflicts can happen due to different versions of the upper ontology used and ontologies that are not fully orthogonal and interoperable. Not all OBO Foundry ontologies use the same version of the BFO and differences based on temporal relations could cause issues in future work. Such challenges can be addressed. Inconsistent high level organization of cell terms were also found during the BCGO development but were solved by coordination with CL, CLO and OBI developers. For example, ‘cell line cell’, has been defined differently in OBI, CL and CLO. After discussion, agreement was reached on the definition of ‘cell line cell’ and its placement in CLO. OBI has deprecated ‘cell line cell’ and imported the term from CLO.

With well-established OBO Foundry reference ontologies and ontology modularity tools, application ontology development can be rapid using this approach through maximal reuse of existing resources, a key principle of ontology development.

## ACKNOWLEDGEMENTS

We thank Dr. E. Greenfest-Allen for her valuable advice. We thank the OBI, CL, and CLO developers, especially Dr. M. Brush, for discussion and consistent implementation of high level cell terms in three ontologies. This research is supported by NIH grant 1R01GM093132-01 and by 5 U01 DK 072473.

